# Limited intra-host diversity and background evolution accompany 40 years of canine parvovirus host adaptation and spread

**DOI:** 10.1101/714683

**Authors:** Ian E.H. Voorhees, Hyunwook Lee, Andrew B. Allison, Robert Lopez-Astacio, Laura B. Goodman, Oyebola O. Oyesola, Olutayo Omobowale, Olusegun Fagbohun, Edward J. Dubovi, Susan L. Hafenstein, Edward C. Holmes, Colin R. Parrish

## Abstract

Canine parvovirus (CPV) is a highly successful pathogen that has sustained pandemic circulation in dogs for more than 40 years. Here, integrating full-genome and deep sequencing analyses, structural information, and *in vitro* experimentation, we describe the macro- and micro-scale features that have accompanied CPV’s evolutionary success. Despite 40 years of viral evolution, all CPV variants are >∼99% identical in nucleotide sequence, with only a limited number (<40) of mutations becoming fixed or widespread during this time. Notably, most changes in the major capsid protein (VP2) are nonsynonymous and fall within, or adjacent to, the overlapping receptor footprint or antigenic regions, suggesting competitive selective pressures have played a key role in CPV evolution and likely constrained its evolutionary trajectory. Moreover, among the limited number of variable sites, CPV genomes exhibit complex patterns of variation that likely include parallel evolution, reversion, and recombination, making phylogenetic inference difficult. Additionally, deep sequencing of viral DNA in original clinical samples collected from dogs and other host species sampled between 1978 and 2018 revealed few sub-consensus single nucleotide variants (SNVs) above ∼0.5%, and experimental passages demonstrate that substantial pre-existing genetic variation is not necessarily required for rapid host receptor driven adaptation. Together, these findings suggest that although CPV is capable of rapid host adaptation, relatively low mutation rate, pleiotropy, and/or a lack of selective challenges since its initial emergence have reduced the long-term genetic diversity accumulation and evolutionary rate. Hence, continuously high levels of inter- and intra-host diversity are not intrinsic to highly adaptable viruses.

**IMPORTANCE:** Rapid mutation rates and correspondingly high levels of standing intra-host diversity and accumulated inter-host diversity over epidemic scales are often cited as key features of viruses with the capacity for emergence and sustained transmission in a new host species. However, most of this information comes from studies of RNA viruses, with relatively little being known about that evolutionary processes that occur for viruses with DNA genomes. Here we provide a unique model of virus evolution, integrating both long-term global-scale and short-term intra-host evolutionary processes of a virus in a new host animal. Our analysis reveals that successful host jumping and sustained onward transmission does not necessarily depend on a high level of intra-host diversity or result in the continued accumulation of high levels of long-term evolution change. These findings indicate that all aspects of a virus’s biology and ecology are relevant when considering their adaptability.

## INTRODUCTION

The emergence of viruses to cause epidemics in new hosts poses a constant threat to humans and other animals, as their appearance in immunologically naïve populations may result in widespread disease and a major health burden. In many instances newly emerging viruses will fix adaptive mutations that increase replication and facilitate onward transmission in the new host species (1,2). However, it is not clear how rapidly phenotypically relevant mutations arise and become selected under different circumstances, how many such changes are needed for host adaptation and sustained transmission, and whether there is a complex series of mutational changes that have to occur in a specific order. Likewise, the relationship between intra-host genetic diversity and long-term inter-host evolution and adaptation remain poorly defined. Clarifying the sources, mechanisms, and extent of viral variation and evolution during known emergence events will therefore provide information central to understanding the risks of similar events in the future.

Canine parvovirus (CPV), a non-enveloped single stranded DNA virus (family *Parvoviridae, genus Protoparvovirus*), is an extremely successful viral pathogen that overcame the evolutionary hurdles associated with cross-species transmission to emerge and spread in multiple host species. The original CPV strain (designated CPV type 2 (CPV-2) to distinguish it from the distantly related minute virus of canines) arose as a variant from a group of closely related viruses circulating among other carnivores, and was first recognized during early 1978 as the causative agent of a disease in dogs characterized by vomiting and diarrhea, especially among young dogs, and myocarditis in neonatal puppies (3–5). By mid 1978 CPV-2 had reached pandemic proportions, and it was replaced by a new genetic variant designated CPV-2a by the end of 1980 (6,7). This new variant contained a group of 5 amino acid changes in the virus capsid protein (VP) gene (7) that enabled it to expand its host range among carnivores, most notably gaining the ability to infect and transmit between domestic cats (8). All CPV variants circulating today are descendants from this original emergence event and remain significant pathogens of dogs, despite the widespread use of effective vaccines (9,10).

A key aspect in understanding the cross-species transmission and continued evolution of CPV and the closely related carnivore parvoviruses is the interaction with their host cell receptor, transferrin receptor type-1 (TfR) (11). Gaining the ability to properly bind the canine TfR for cell infection was a critical evolutionary step for CPV (12,13), and comparisons of FPV and CPV sequences revealed that the canine host range was determined by 3 or 4 mutational differences within a small region of the VP2 capsid surface and involving at least 3 surface loops contributed by 2 symmetrically arranged VP2 molecules (14–16). These capsid mutations overcame a biochemical canine host range barrier - the presence of an N-linked glycosylation site associated with residue 384 of the canine TfR - that blocked receptor binding by the capsids of the ancestral-like viruses from cats, mink and raccoons (12,17). Mutations that arose later in the evolution of CPV, particularly those that gave rise to the CPV-2a variant, may have evolved in other hosts besides dogs as changes of the CPV-2a-specific codons to other residues have been found in viruses from raccoons, foxes, and mink (18,19). Such mutations likely reflect adaptation to the viruses overcoming host barriers to infection or optimizing their interactions with TfR binding sites (19). Recently, the TfR-CPV interaction has been further defined using cryo-electron microscopy (cryoEM) (Lee, H., et al. personal communication), providing atomic resolution structural information of the viral capsid-host cell receptor interface.

Other important interactions at the virus-host interface include antibody binding, and selective pressures for antigenic variation has likely played a role in driving the ongoing evolution and success of these viruses in nature (7,20). Studies defining the binding sites of 8 different mouse or rat monoclonal antibody (MAb) antigen binding fragments (Fabs) revealed that over 65% of the accessible viral capsid surface was covered by antibody footprints, although the mutations that controlled natural or experimentally selected antigenic variation fell within two relatively small regions (21,22). Importantly, the TfR binding site overlaps with some key MAb binding sites, and many capsid mutations may alter both antibody and TfR binding, such that there are likely complex relationships between the host selection by the TfR and antigenic selection by antibody immunity (12,23).

While many studies have identified specific consensus-level host-adaptive mutations of the major viral capsid, none have sought to survey the full suite and distribution of mutations that have become fixed or widespread during CPV’s emergence in dogs and subsequent 40 years of circulation. Moreover, the link between intra-host evolution and that at the epidemiological scale remains unclear. In particular, is CPV’s capacity to emerge in multiple hosts caused by high levels of standing genetic variation within individual hosts? The ability to perform genome-wide deep sequencing on natural and experimental viral samples now allows us to more completely define the variability and evolution within the viral population. Other studies on emerging viruses have primarily focused on RNA viruses and have revealed that individual hosts can harbor viruses with multiple SNVs [reviewed in (24)], the consequence of highly error-prone replication (25,26). Far less is known about the extent and properties of sequence variation within populations of DNA viruses, including ssDNA viruses such as the parvoviruses. However, despite being replicated by DNA polymerases that are very high-fidelity when replicating the host cell genome, CPV evolved relatively rapidly in nature during its initial emergence, with evolutionary rates in the range of ∼10^-4^ substitutions/site/year (27,28), and the virus is able to rapidly adapt to both variant host receptors and to antibody selection in tissue culture (19,22).

Herein, we describe the emergence and sustained transmission of CPV over 40 years across multiple evolutionary scales. We combine genome-wide and deep sequence analyses of a collection of CPV-related viruses in original clinical samples from dogs and alternative host species collected in 1978 and during the subsequent 40 years of virus evolution, as well as experimental studies of host adaptation to determine (i) the full suite of genetic changes that have become fixed or widespread during emergence and pandemic circulation of CPV and (ii) the extent and structure of intra-host genetic diversity that underpins this long-term evolution. These data, informed by recently resolved high-resolution molecular structures, provide a model encompassing both the macro- and micro-scale features enabling a virus to emerge and sustain pandemic spread in a new host.

## RESULTS

The viruses sequenced as part of this study represent a record spanning the entire 41 years of evolution of CPV in dogs, as well as that occurring in the related viruses recovered from raccoons, arctic foxes, and raccoon dogs. Most of these samples originated from the United States, but samples from the Australia, Nigeria, France, Germany and Finland were also included. A total of viral 46 genomes from original fecal or intestinal material were sequenced, of which 35 were deep sequenced (Table 1). Viral genomes from various tissues of an acutely infected puppy, as well as from experimentally passaged viruses in different host cells in culture were also included in our analyses.

**Table 1.** Naturally circulating viruses sequenced in this study. Accessions to be assigned are listed as “TBA”. SRA datasets for deep sequenced samples only.

| Isolate name | Collection year | Country | Host | Accession (consensus) | SRA accession (reads) |
| --- | --- | --- | --- | --- | --- |
| CPV12 | 1978 | United States | Dog | TBA | TBA |
| CPV6 | 1979 | United States | Dog | TBA | TBA |
| CPV9 | 1979 | United States | Dog | TBA | TBA |
| CPV5* | 1980 | United States | Dog | EU659116.1 | TBA |
| CPV63 | 1980 | United States | Dog | TBA | TBA |
| RDPV122 | 1980 | Finland | Raccoon Dog | TBA | TBA |
| RDPV123 | 1980 | Finland | Raccoon Dog | TBA | TBA |
| CPV13* | 1981 | United States | Dog | EU659118 | TBA |
| CPV81 | 1982 | Australia | Dog | TBA | TBA |
| BFPV2 | 1983 | Finland | Blue Fox | TBA | TBA |
| CPV27 | 1983 | United States | Dog | TBA | TBA |
| CPV29 | 1983 | United States | Dog | TBA | TBA |
| CPV30 | 1983 | United States | Dog | TBA | TBA |
| CPV31 | 1983 | United States | Dog | TBA | TBA |
| CPV33 | 1983 | United States | Dog | TBA | TBA |
| CPV58 | 1983 | France | Dog | TBA | TBA |
| CPV35 | 1984 | United States | Dog | TBA | TBA |
| CPV36 | 1984 | United States | Dog | TBA | TBA |
| CPV39 | 1984 | United States | Dog | TBA | TBA |
| CPV54 | 1984 | France | Dog | TBA | TBA |
| CPV48 | 1985 | United States | Dog | TBA | TBA |
| CPV87 | 1985 | Australia | Dog | TBA | TBA |
| CPV50 | 1986 | United States | Dog | TBA | TBA |
| RDPV124 | 1986 | Finland | Dog | TBA | TBA |
| CPV220 | 1993 | United States | Dog | TBA | N/A |
| CPV305 | 1993 | United States | Dog | TBA | TBA |
| CPV307 | 1993 | United States | Red Wolf | TBA | TBA |
| CPV353 | 1996 | United States | Dog | TBA | TBA |
| CPV604 | 2007 | United States | Dog | TBA | TBA |
| RACCPV09 | 2009 | United States | Raccoon | TBA | TBA |
| RacCPV2 | 2011 | United States | Raccoon | TBA | TBA |
| CPV617 | 2011 | United States | Dog | TBA | TBA |
| CPV606 | 2014 | United States | Dog | TBA | TBA |
| CPV601 | 2016 | United States | Dog | TBA | TBA |
| CPV603 | 2017 | United States | Dog | TBA | TBA |
| CPV607 | 2018 | Nigeria | Dog | TBA | N/A |
| CPV608 | 2018 | Nigeria | Dog | TBA | N/A |
| CPV609 | 2018 | Nigeria | Dog | TBA | N/A |
| CPV610 | 2018 | Nigeria | Dog | TBA | N/A |
| CPV614 | 2018 | Nigeria | Dog | TBA | N/A |
| CPV615 | 2018 | Nigeria | Dog | TBA | N/A |
| CPV616 | 2018 | Nigeria | Dog | TBA | N/A |
| CPV605 | 2018 | Nigeria | Dog | TBA | TBA |
| CPV611 | 2019 | United States | Dog | TBA | N/A |
| CPV612 | 2019 | United States | Dog | TBA | N/A |
| CPV613 | 2019 | United States | Dog | TBA | N/A |

Sequencing provided nearly complete genome coverage, spanning all major reading frames and lacking only the distal portions of the 5’ and 3’ terminal untranslated regions (UTRs) (see Materials and Methods and Fig. S1). For samples that were deep sequenced in order to identify sub consensus single nucleotide variants (SNVs), ∼5000-fold coverage per site was achieved after read processing and coverage normalization. A short region (∼150nts in length) flanked by a poly A and a G-rich sequence region near the start of the VP2 gene did not sequence or assemble efficiently in any of the natural or experimental samples or control plasmids. While the low coverage in this region (the “low coverage region” (LCR) in Figs. 4-6) did not affect consensus calls, it did affect our ability to accurately call SNV frequencies and was therefore excluded when measuring intra-host diversity. To confirm and define the sensitivity of our sub-consensus variant calling in the rest of the CPV genome, artificially created heterogenous virus populations were created by serially diluting two different infectious clones that differed at 39 nucleotide positions (i.e. 39 SNVs). These control samples, which were processed alongside the natural samples, provided evidence that our methods could consistently detect SNVs at very low proportions (<1%) across the genome (Fig. S2). We therefore set a cut-off of SNV detection of 0.5%. For all analyses in this study genome nucleotide numbering starts at the first position in the NS1/2 gene coding region, while amino acid numbering begins at the first methionine for each protein.

### Limited background evolution of CPV during 40 years of viral circulation

New CPV consensus sequences determined for the viruses analyzed here were combined with all other CPV sequences isolated from dogs and raccoons spanning both the NS and VP coding regions (hereafter referred to as the “full-genome”) obtained from GenBank. CPV sequences derived from raccoons were also included as these hosts have been suggested to play a major role in the long-term evolution of these viruses (18). This data set (222 genomes) was used to survey the distribution and nature of mutations that have arisen and become fixed or widespread in all open reading frames and introns of the CPV during its circulation (Table 2, Fig. 1). We included all of those mutations that differed from the earliest CPV full-genome sequence in the database (“CPV strain N” M19296.1) and were present in >10% (arbitrarily chosen) of the sequences analyzed. Overall, 17 synonymous and 22 non-synonymous mutations met these criteria. Of the 17 synonymous mutations, 10 fell within the non-structural gene region of the genome, 3 in the VP1 intron, and 3 within the VP coding regions. Of the non-synonymous changes, 4 were found in NS1, 4 in NS2, 1 was exclusive to VP1, 11 were in the VP1/2 gene overlapping region, and 2 were in the small alternative open reading frame found near the N-terminus of VP2, the SAT protein gene (Fig. 1).

**Figure 1.**
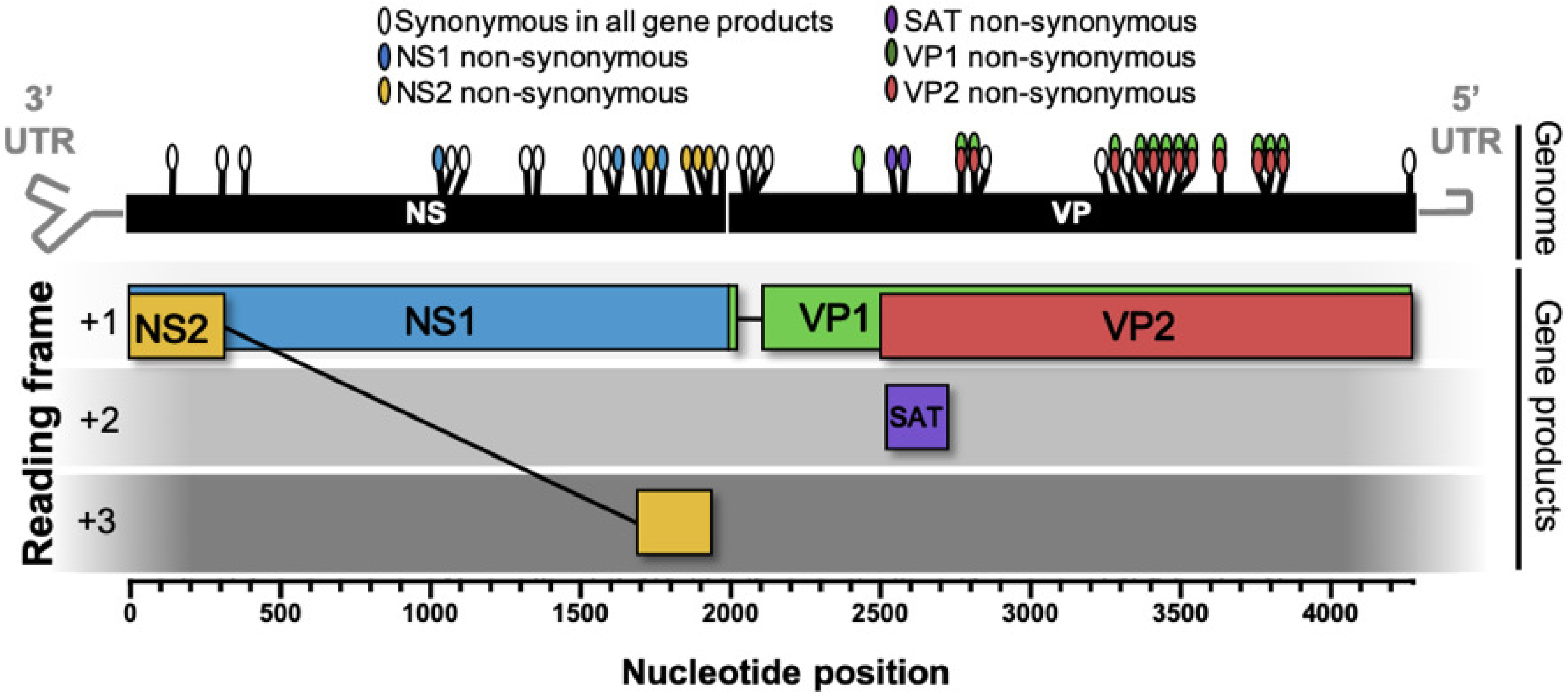
Survey of mutations in the CPV genome that have arisen and become widespread or fixed since the virus first emerged in dogs in 1978. Schematic diagram of the CPV genome and gene products with locations and coding effect (colored ovals) of substitutions. The analysis was based on 225 full-genome CPV sequences from virus isolated in dogs or raccoons. Genome region of analysis includes all coding regions as well as the VP1 intron (black boxes). Nucleotide position numbering begins at NS1/2 translational start site. Grey hairpins in genome represent terminal untranslated regions (UTRS) which were not included in this analysis.

**Table 2.** Position and translation effect of the CPV substitutions depicted in Fig. 1. Translation effects are colored as in Fig.1. Nucleotide numbering begins at the first coding site of the NS1/2 gene. Amino acid numbering is based on each respective gene translational start site. VP2 position is given for substitutions within the VP1/VP2 overlapping ORFs. Asterisk (*)indicates positions that arose during the CPV-2 to CPV-2a global sweep.

| NT Position | NT Change | Translation Effect |
| --- | --- | --- |
| 177 | A->G | Synonymous |
| 342 | T->C | Synonymous |
| 411 | G->A | Synonymous |
| 1053 | T->A | NS1 N351K |
| 1062 | A->G | Synonymous |
| 1098 | A->G | Synonymous |
| 1353 | A->G | Synonymous |
| 1377 | T->C | Synonymous |
| 1542 | T->C | Synonymous |
| 1629 | T->C | Synonymous |
| 1631 | A->T | NS1 Y544F |
| 1714 | G->A | NS1 E572K |
| 1752 | A->G | NS2 T94A |
| 1790 | T->C | NS1 L597P, |
| 1875 | A-> G | NS2 T135A |
| 1923 | G->A | NS2 D151N |
| 1926 | A->G | NS2 M152V |
| 1975 | T->C | Synonymous |
| 2063 | A->G | Intron |
| 2085 | A->G | Intron |

| NT Position | NTChange | Translation Effect |
| --- | --- | --- |
| 2086 | A->G | Intron |
| 2432 | A->G | VP1 K116R |
| 2550 | A->G | SAT N10S |
| 2574 | T->A | SAT L18Q |
| 2773 | A->T | VP2 M87L* |
| 2816 | T->C | VP2 I100T* |
| 2817 | T->C | Synonymous |
| 3246 | T->C | Synonymous |
| 3314 | T->A | VP2 F267Y |
| 3345 | T->C | Synonymous |
| 3403 | T->G | VP2 S297A |
| 3413 | C->G | VP2 A300G* |
| 3427 | G->T | VP2 D305Y* |
| 3484-3485 | TA->CT | VP2 Y324I |
| 3637 | A->G | VP2 N375D* |
| 3790 | A->G | VP2 N426D |
| 3792 | T->A | VP2 D426E |
| 3832 | A->G | VP2 T440A |
| 4266 | T->C | Synonymous |

\* Indicates mutations that differentiate CPV-2a from CPV2

Recently determined structures showing the footprint of the TfR from a black backed jackal, or of antibody Fab-capsid co-structures were used to characterize the structural context of the capsid non-synonymous substitutions identified in our survey (Fig. 2). Of the non-synonymous mutations in VP2, 3 (amino acid positions 87, 305, and 300) fell directly within the footprint of the TfR or were immediately adjacent to that footprint, and those also fell within or immediately adjacent to defined MAb Fab footprint regions (Fig. 2). All remaining naturally variable capsid surface residues fell within or immediately adjacent to the monoclonal antibody (MAb) antigen binding fragment (Fab) footprints, but did not overlap the TfR footprint (Fig.2). Additionally, 3 naturally occurring non-synonymous mutations in the capsid (VP2 residues 87, 101, and 375) were found located beneath the capsid surface, perhaps resulting in indirect and relatively subtle capsid changes. The overlap of receptor and MAb Fab footprints and limited number of mutations on the surface suggests that the CPV capsid is likely evolutionary constrained by competing selective pressures of TfR and antibody interactions.

**Figure 2.**
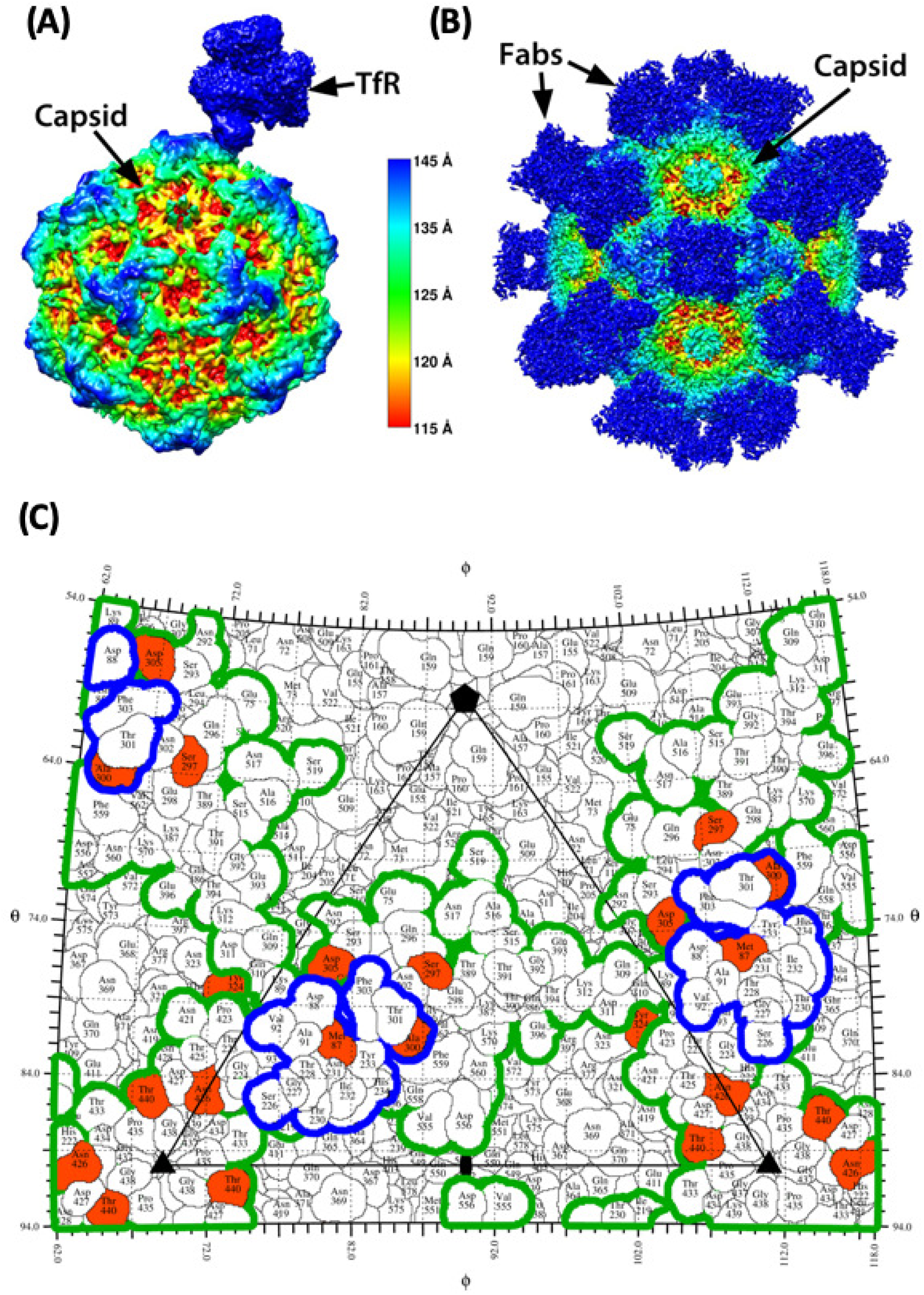
Structural context of natural CPV variants and their relation to TfR and MAb Fab footprints. High resolution structure of the transferrin receptor (A) and monoclonal antibody Fabs (B) bound to the CPV capsid colored by capsid radial distance identifies the sites of contact. (C) One asymmetric unit of CPV capsid, showing the footprint of the TfR (blue outline) and the combined footprints of 8 different antibodies (green outline). Red residues - varied during evolution of CPV in dogs. All structures determined by cryoEM. See main text for references.

Less is known about the possible phenotypic effects of the mutations that occurred within the viral non-structural genes encoding for NS1, NS2, and the small alternatively translated protein (SAT) contained within VP2. However, all of these non-structural proteins interact with cellular proteins and processes as demonstrated in CPV or other closely related parvoviruses (29–36), and different selective pressures may therefore exist within different host species. In NS1, most non-synonymous mutations fell within the C-terminal region, outside of the nickase and the helicase regions as defined in closely related parvoviruses (37,38), which are conserved in structure, amino acid sequence, and function. Further experimental mutagenesis is needed to understand the phenotypic effects of naturally occurring changes in the non-structural genes of these viruses.

### Phylogenic analyses reveal complex patterns of inter-host evolution and recombination

To determine the evolutionary patterns of CPV at the epidemiological scale we performed full-genome phylogenetic analyses. All CPV full-genome sequences isolated from dogs and raccoons and raccoon dogs were included in this analysis.

Screening for recombination in these data using the Recombination Detection Program (RDP4; default settings) (39) revealed a common topological break point situated within the VP1 exclusive coding region. However, as most of the putative recombination events received limited bootstrap support in the phylogenetic analysis, it is difficult to conclusively determine if this breakpoint results from recombination, differential selective pressures on the NS and VP genes, or a combination of both factors. For example, 6 potential distinct recombination events were identified accounting for a total of 26 of the 225 genomes analyzed. However, there were discrepancies in determining the recombinants between the various detection methods, with only 4 potential events, accounting for only 6 genomes in total, being supported by 2 or more detection methods. Of these 6 genomes, 3 were recent isolates from Nigeria sequenced as part of this study (CPV610, CPV614, and CPV615), with NS region of these sequences resembling that of a 2014 Chinese isolate (accession number KT382542) but an unknown VP region parent. The remaining 3 genomes were database sequences (accession numbers KM457139.1, KX774252, and KR002800), only 1 of which (KM457139.1) had been described previously as a recombinant (40). For the purposes of this study, the 6 genomes supported by 2 or more detection methods with the default RDP4 analysis, were removed from our phylogenetic analysis.

A maximum likelihood tree was then inferred from the remaining 219 full-genome sequences, and annotated by country of sample origin as well as by the mutations of interest (Figs. 3). This analysis clearly identified the 5 non-synonymous changes associated with the rise and pandemic spread of CPV-2a viruses in the early 1980s (Fig. 3A) (6,41). However, after this evolutionary leap, support for phylogenic structure greatly diminishes, and the evolutionary topology was marked by a distinct lack of resolution. This poor phylogenic support for the specific evolutionary analysis is often overlooked in studies of CPV natural evolution, or avoided by subsampling select sequences. While some small isolated clades of high bootstrap support are apparent, such as a clade of recent Chinese viruses, a single Italian isolate of Chinese origin, and several Nigerian viruses, most sequences sampled appear to be minor variants of a common “pan genome” template. In addition, many of the mutations acquired are short-lived, in that they fall on tip branches, and both parallel evolution and reversion appear to be common occurrences. For example, VP2 residue 426 is an Asn in FPV-like viruses, and in CPV-2 and CPV-2a genotypes, but antigenically distinct CPVs resulting from mutations at this position to Asp or Glu have been named CPV-2b and CPV-2c, respectively. However, these amino acid changes do not appear to segregate into strongly supported monophyletic groups of viruses and likely arose multiple times after the emergence of CPV-2a (Fig. 3B). Other such mutations in the capsid that have potential phenotypic impacts based on their proximity to the TfR MAb Fab footprints include Ala at VP2 residue 297 Glu at VP2 residue 440. While it is possible that frequent recombination could also produce these patterns, with a low numbers of true segregating sites and, likewise, a low baseline bootstrap support, it is difficult to estimate the relative contributions of parallel evolution, reversion, and recombination on the observed genetic homoplasy.

**Figure 3.**
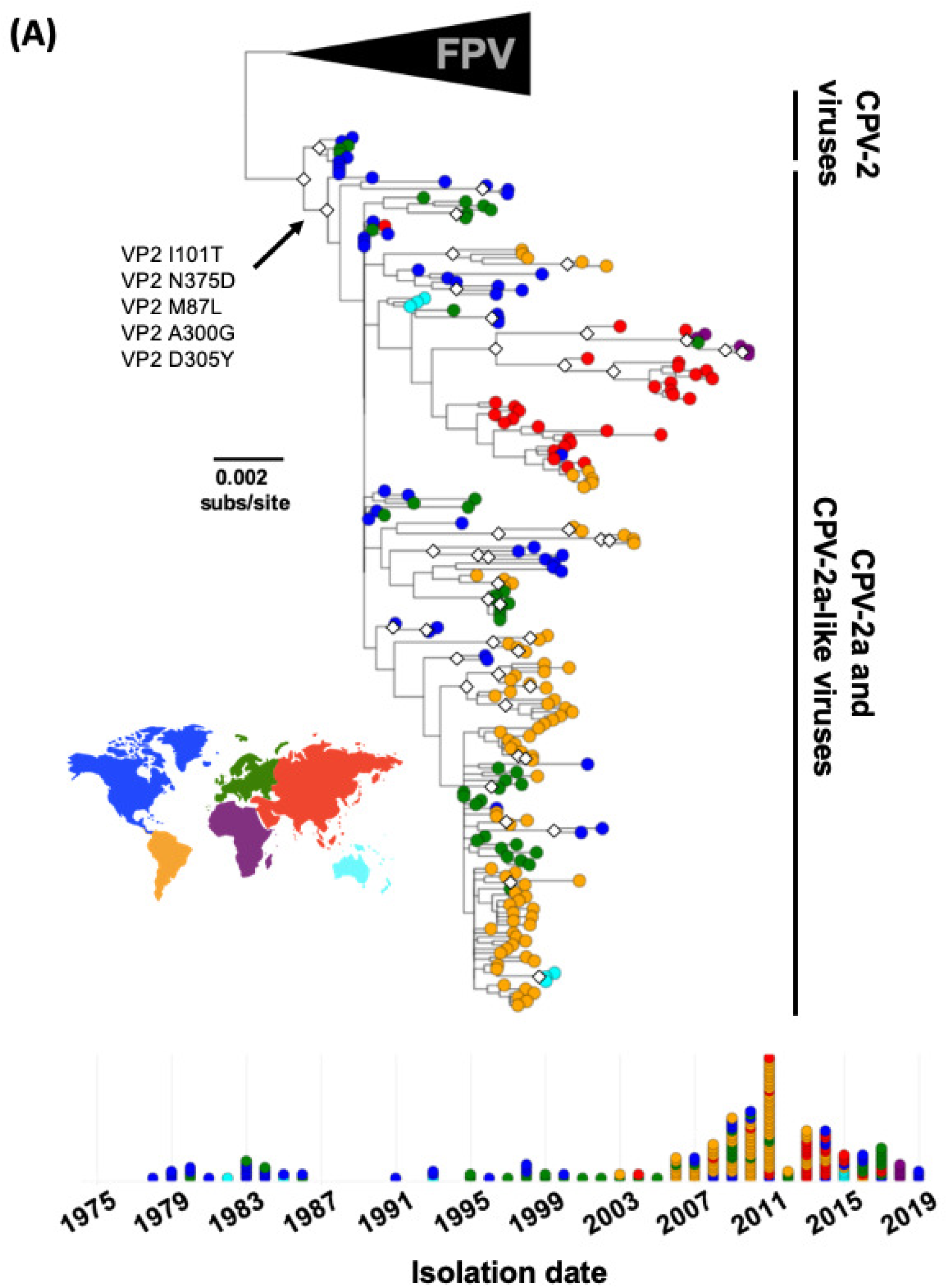
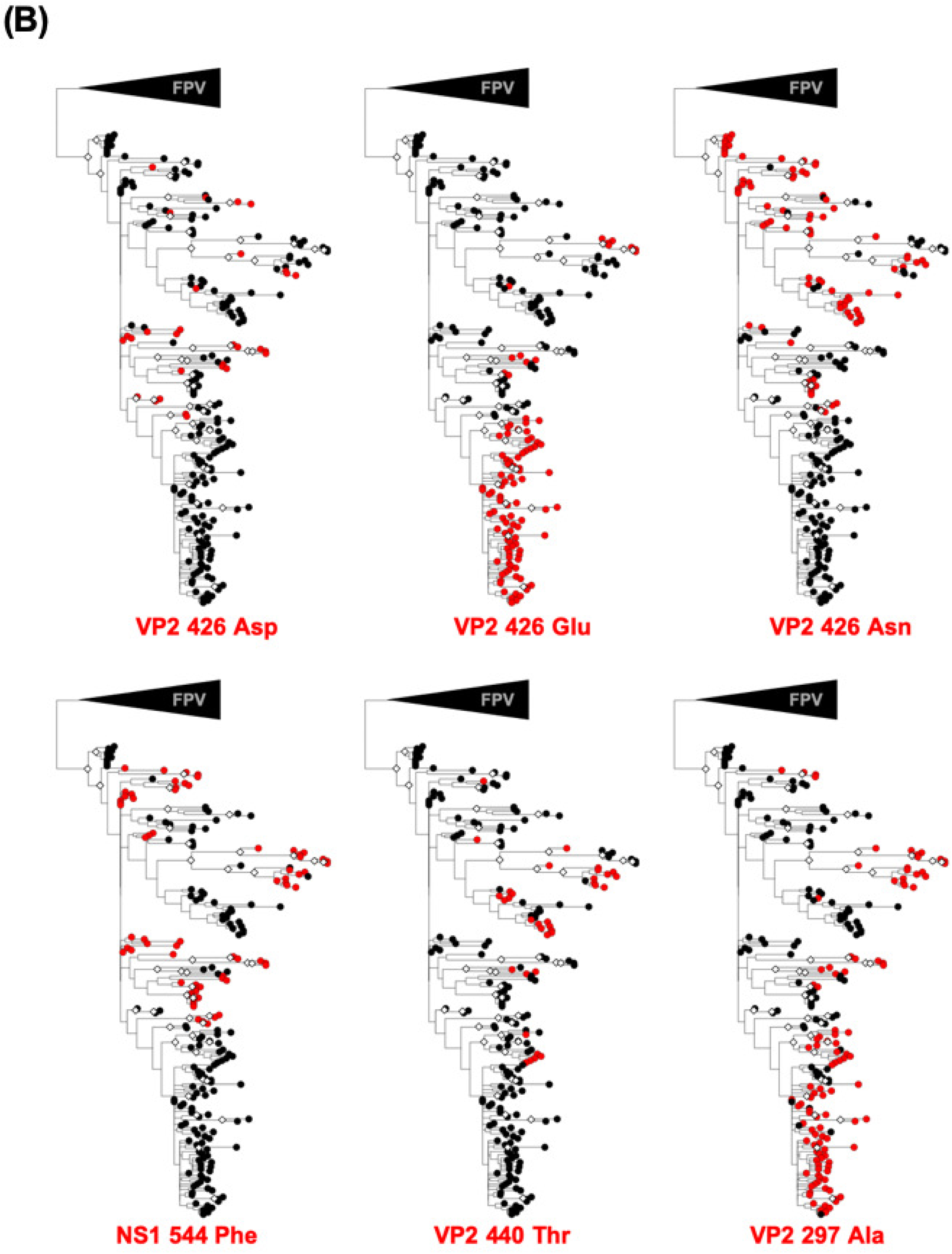
Evolutionary relationships among CPV full-genomes. Sequences analyzed include all complete CPV genomes available that were isolated from dogs and raccoons, and raccoon dogs. An FPV-like virus clade was used as an outgroup and bootstrap values >75% are indicated with white diamond (◇) node. (A) Phylogeny with tip shapes colored by geographic origin (continent). The branch and mutations leading to the emergence and pandemic replacement CPV-2a viruses are indicated. A timeline of samples (lower panel) shows the temporal spread of sequences analyzed. (B) Phylogenies annotated by mutations of interest that appear to have multiple evolutionary origins.

### Extremely low levels of intra-host diversity among CPVs circulating in dogs during 40 years of virus evolution

The natural virus samples collected from dogs showed few sub-consensus SNVs within the deep sequences of any viral DNA collected from infected animals at any time during the CPV pandemic (Fig. 4). Importantly, several deeply sequence samples were collected in the late 1970s and early 1980s, around the time of the global spread of CPV-2, and its replacement by CPV-2a. Sequences from this time are underrepresented and could shed light on this important and well documented evolutionary transition. Among the 28 natural clinical samples sequenced from dogs (>150,000 bases covered), we observed a total only 8 sub-consensus SNVs above 0.5%. Of these SNVs, only 2 (resulting in non-synonymous changes of VP2 D375N and VP2 I555V) were also present as mutations in the consensus sequence analysis (see Table 2, Fig. 1), and both were found at frequencies below <2% in a single viral sample (CPV48) collected in 1985 in the USA. In addition, no variant sequences or sub-consensus SNVs above 0.5% were observed in viral DNA from different tissues of a naturally infected dog (Fig. 5), suggesting that tissue-specific adaptation of the virus is not a major feature of this virus in dogs, at least in this case. Individual coverage and variant call plots for all samples collected from dogs provided in Figs. S3A-C.

**Figure 4.**
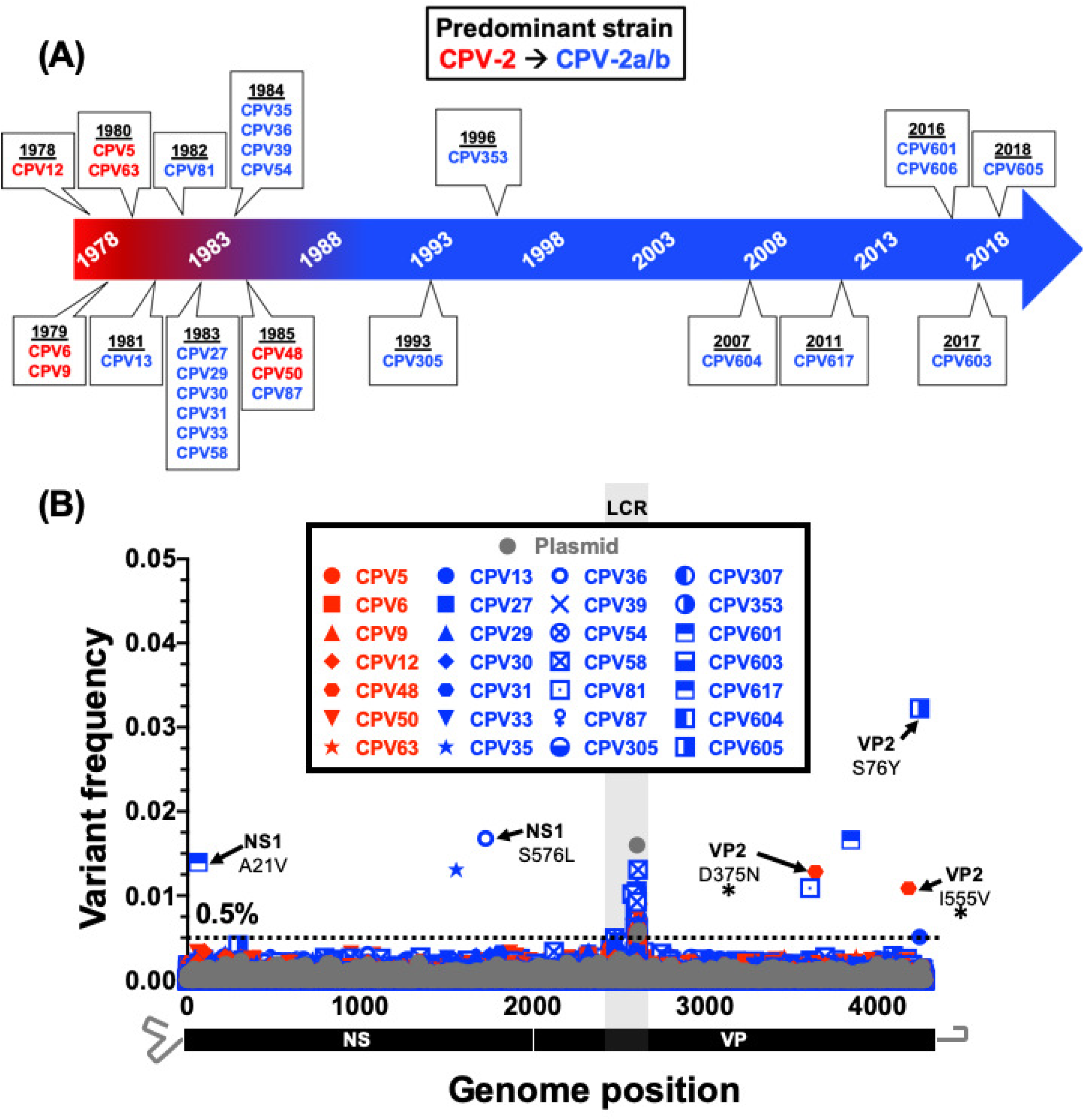
Intra-host viral diversity among dogs infected with CPV-2 or CPV-2a sampled during 40 years of circulation. (A) Time-line displaying dates of CPV from original fecal material samples that were deep sequenced as part of this study. The dominant circulating strain and of each isolate are indicated by red and blue time-line shading and text, respectively. (B) Position and frequency of all dominant sub-consensus SNVs are displayed. Red and blue symbol coloring as in (A). SNVs resulting in non-synonymous changes are indicated. The non-synonymous changes also found in survey of natural samples (see Fig. 1) are indicated with an asterisk (*). Deep-sequenced plasmid (in grey) provides a baseline sequencing error rate and the low coverage region (LCR) is indicated with grey shading and positions in this region were omitted from variant calling analyses.

**Figure 5.**
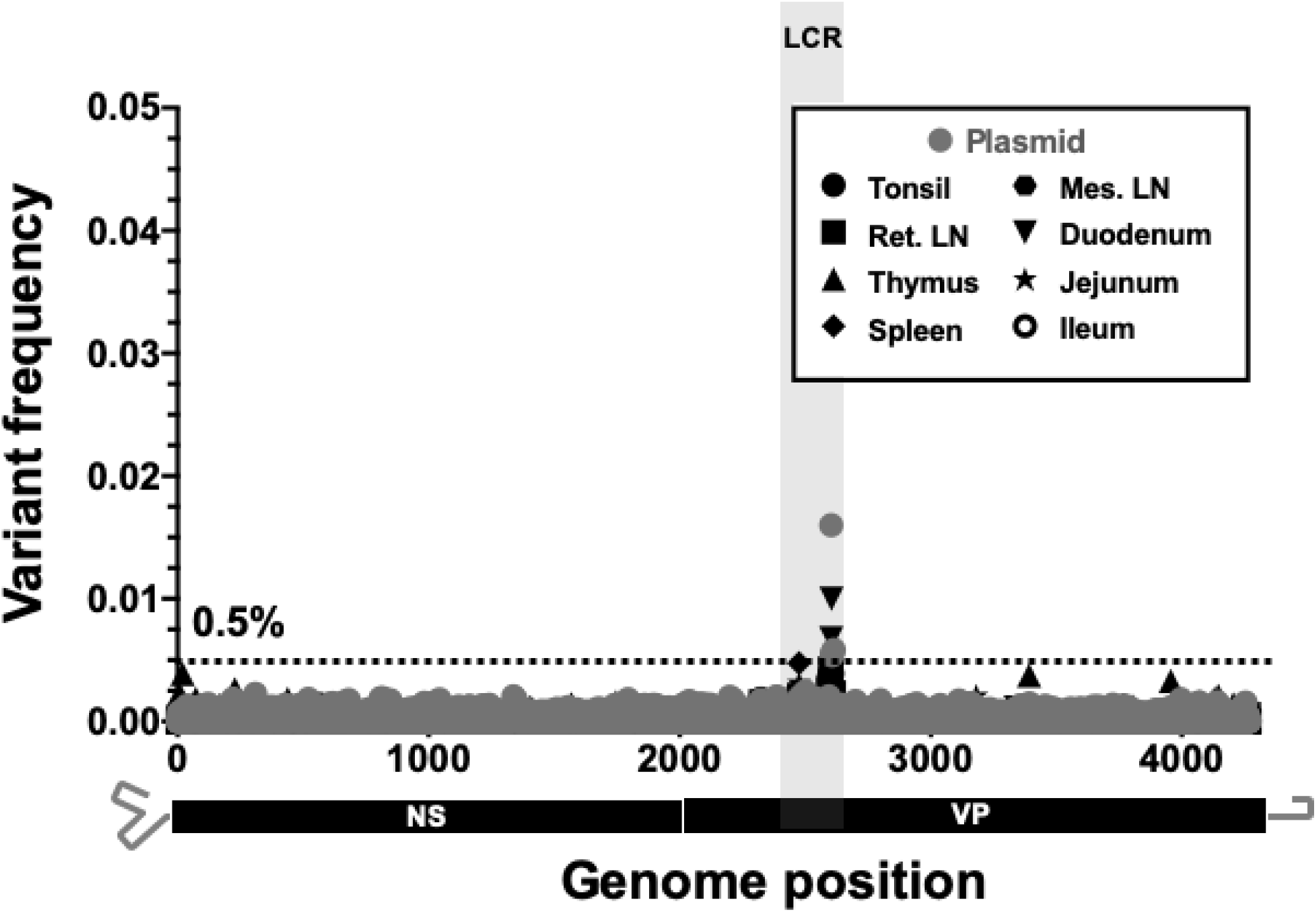
Intra-host viral diversity in different tissues of a single dog during an acute CPV-2a infection. Position and frequency of all dominant sub-consensus SNVs are displayed. Deep-sequenced plasmid (in grey) provides a baseline sequencing error rate and the low coverage region (LCR) is indicated with grey shading and positions in this region were omitted from variant calling analyses.

### Extremely low levels of intra-host diversity among CPVs from other host species

While CPV has spread and been maintained in the large domestic dog population of the world, CPV-related viruses infecting other hosts often show specific changes in the capsids which are likely associated with adaptation to variation of the specific TfRs in those animals (15,19). Notably, our deep sequencing of 5 CPVs and 2 FPV intra-host populations from raccoons (*Procyon lotor*) in the USA, raccoon dogs (*Nyctereutes procyonoides*) in Finland, a red wolf (*Canis lupus rufu*s) in the USA, and an Arctic (blue) fox (*Vulpes lagopus*) in Finland also revealed low levels of intra-host diversity similar to those seen in the canine samples, with a total of only 3 sub-consensus SNVs detected above ∼ 0.5% (Fig. 6). All 3 of these sub-consensus SNVs were non-synonymous in at least one gene product, but none were at positions known to be involved with host adaptation. Individual coverage and variant call plots for all samples collected from non-dog hosts are provided in Fig. S3D.

**Figure 6.**
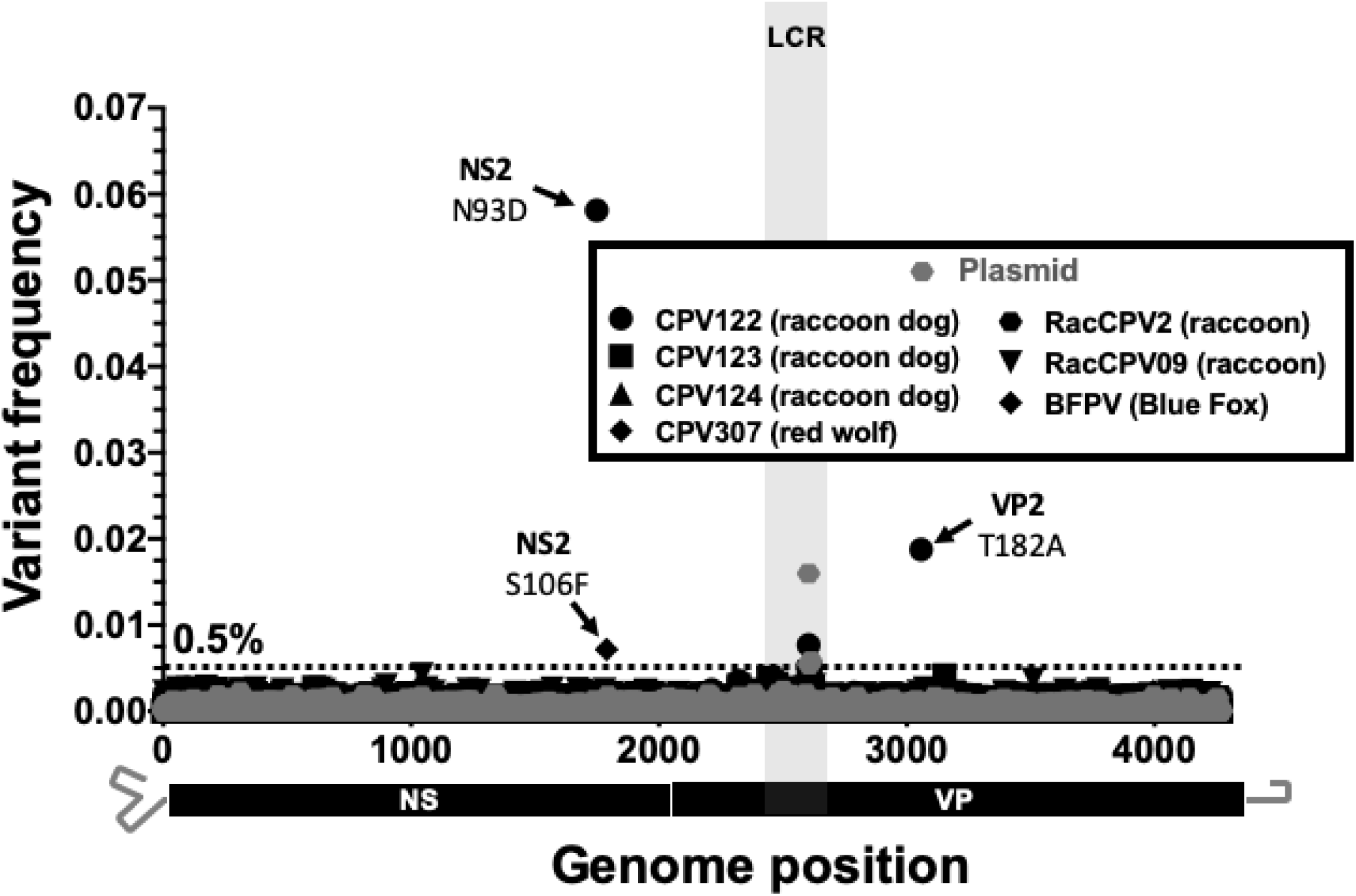
**Intra-host viral diversity among alternative host species infected CPV-2, CPV-2a, or FPV.** Position and frequency of all dominant SNVs are displayed. Sequenced viruses include CPV-2 isolates from raccoon dogs (CPV122-124), CPV-2a isolates from a red wolf (CPV307) and a raccoon (RACCPV09), and FPV isolates from an arctic blue fox (BFPV) and Raccoon (RacCPV2). Deep-sequenced plasmid (in grey) provides a baseline sequencing error rate and the low coverage region (LCR) is indicated with grey shading and positions in this region were omitted from variant calling analyses.

### Substantial intra-host diversity is not a prerequisite for rapid host receptor-driven adaptation in cell culture

To examine the selective dynamics of the host-specific mutations in these viruses we passaged a CPV-2a-derived virus in cells from a domestic cat (*Felis catus*) and from a gray fox (*Urocyon cinereoargenteus*). Those host cells have previously been shown to select for mutations in CPV-related viruses from different sources (15,19), in studies that primarily followed consensus mutations using Sanger sequencing. The virus used was a 1984 isolate that contained the change of VP2 residue 426 to Asp (sequence “CPV39” in this study) (6,20), and an infectious stock was prepared from an infectious plasmid clone by three passages in feline cells in culture (42). Importantly, no sub-consensus SNVs sites were detected in the plasmid, and very little (all below ∼0.5% of the reads) in the virus stock (Fig.7A), so the starting populations for these experimental viruses resembled those observed in natural samples.

**Figure 7.**
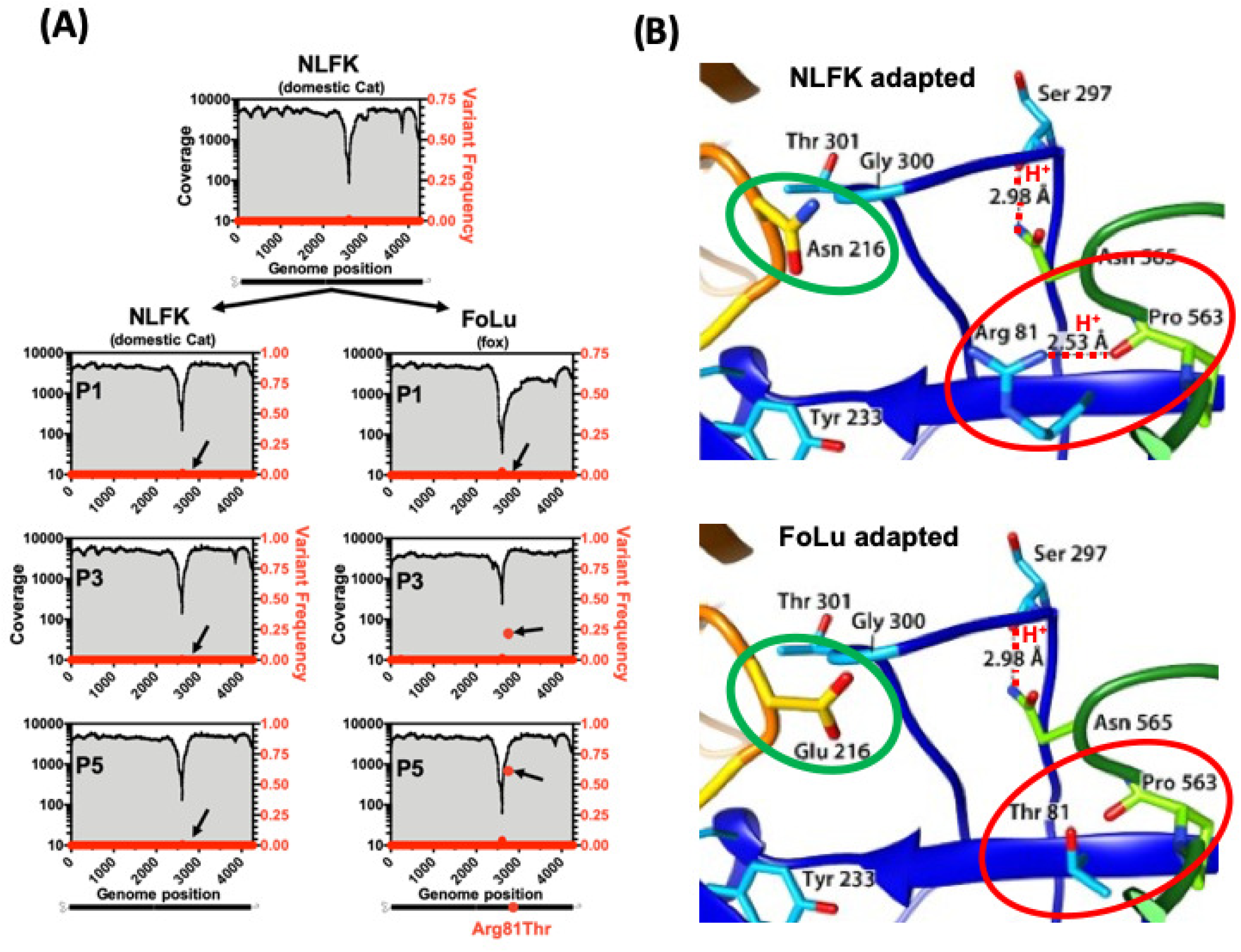
**Receptor-driven adaptation of CPV-2a during a simulated cross-species transmission event.**(A) CPV-2a stock virus originating from transfection of NLFK cells with infectious plasmid (p440, sequence CPV39 in this study) was passaged simultaneously on NLFK and FoLu cell lines and deep sequenced at passage 1, 3, and 5 (P1-5). Coverage (black lines and grey shading) and dominant SNV frequencies (red dots) are displayed. A single non-synonymous substitution (G→C) at nucleotide position 2756 (black arrows) resulting in a VP2 R81T change reached majority consensus by P5 in FoLu cells only. (B) Comparison of cat (upper panel) and fox (lower panel) transferrin receptor (TfR) apical domain amino acid sequences and interacting viral capsid sequences. Residues that differentiate cat and fox TfRs are circled in green. The fox-selected capsid substitution (VP2 R81T), circled in red, results in the loss of a stabilizing hydrogen bond (red dotted line).

After five additional passages of this virus in feline cells, no SNVs above ∼0.5% were observed. However, after three passages in fox cells a single mutation of the codon for VP2 residue 81 from Arg to Thr rose to 20% of the reads, and that represented 60% of the sequence reads (and hence in the majority consensus sequence) after two additional passages in those cells (Fig. 7A). This same VP2 Arg81Thr mutation has been previously described (15) and falls within a region of the capsid structure that stabilizes a loop containing residues 300 and 301, which interact with the TfR (Fig. 7B). By removing a hydrogen bond interacting with this loop, the VP2 Arg81Thr substation likely permits greater flexibility of the capsid. This may allow improved binding or infection through the fox TfR, which differs from the cat in a number of key residues in the two loops of the receptor apical domain that interact with the viral capsid, most notably at position 216 (Fig. 7B, green circles).

## DISCUSSION

CPV first appeared in dogs in the mid-1970s, rapidly emerged to reach pandemic proportions within a few years, and has now maintained global circulation for over 40 years, despite widespread use of effective vaccines and other control measures. While there have been many studies published on the sequence variation of various regions of the CPV genome (recent e.g. (43,43,44)), largely considering the capsid protein genes, here we specifically focused on the extent and pattern of intra-host genomic variation and its connections to long-term virus evolution and the sources of functionally selected mutations. We show that the success of the virus in its new host was accomplished with only a limited number of long-term sustained evolutionary changes and with infections containing little intra-host genetic diversity.

After the initial emergence of CPV-2, and then CPV-2a, there is no evidence of global sweeps of new CPV genotypes, and the evolutionary (i.e. phylogenetic) history of these viruses becomes much more obscure, reflected by low bootstrap supports for most nodes and the limited accumulation of mutation variation despite 40 years of evolution. This lack of accumulated inter-host diversity may mean that most the host adaptation in CPV was achieved early in its evolutionary history in dogs. While some later lineages have accumulated seemingly fixed mutational changes, there are several positions in the CPV genome that appear to have experienced recurrent parallel evolution or reversion, and thus are variable both within and among different clades. Many of the variable sites in the capsid protein gene may be associated with antibody escape and receptor interaction, or both, and it is possible that at least some of these mutations facilitate the fine-tuning of the virus to the host in question, although this will requires careful experimental verification.

There are a number of possible underlying explanations for the relatively limited accumulation of CPV inter-host diversity over 40 years. The simplest is that mutations accumulate at a rate dependent on the background mutation rate of the virus, which is itself lower than the rates seen in RNA viruses (45). This effect may be compounded by the compact nature of the CPV genome (Fig. 1) and the conservation of secondary structures and/or capsid interacting sequences within the ssDNA viral genome (46). In this model most synonymous mutations within the CPV genome would impact multiple functions (i.e. pleiotropy) and hence reduce viral fitness. Similarly, the extensive overlap of the TfR and Fab footprints means that mutations changing the structures of these capsid regions would likely effect both functions. It is therefore possible that only a few sites in the genome can accommodate neutral mutations. This model of low mutation rates and/or extensive pleiotropy is supported by the finding of very low levels of intra-host variation in CPV.

Consistent with the observed low levels of accumulated long-term background evolution over 40 years, our deep-sequencing of natural and experimental viral populations show that high intra-host diversity is not a common feature of these viruses, nor is it a specific barrier to rapid host adaptation in experimental systems (15,19). While replication of the parvoviral DNA and other similar ssDNA viruses by the host cell DNA polymerases appears to be of lower fidelity than that seen for the replication of the host DNA, it is still of higher fidelity than that of RNA viruses (28). This difference between RNA and DNA virus error rates is supported by the lack of intra-host diversity observed in our samples. Similarly, our results also clearly show that co-infection is not a common feature among the CPV samples collected from either dogs or other hosts over the 40 years of sampling, which differs from the results of other studies reporting co-infections between CPV strains or with FPV reported in dogs or cats (40,47–49). However, the finding of recombinant CPV genomes does indicate that co-infections occur on occasion in nature (40,50–52). Thus, while mixed infections appear not to be a predominant feature of CPV infections, they may still have important roles in long-term virus evolution.

Despite these evolutionary constraints associated with relatively low mutation rates and limited intra-host diversity, CPV can still rapidly adapt to acquire new host ranges and evade host humoral immunity - likely by relying on strong and direct selection of relatively small numbers of mutations. For example, during adaptation to alternative host TfRs, single changes in the CPV capsid structure may directly alter the interaction of the capsid with the TfR apical domain, or may alter the flexibility of the receptor-interacting surface of the capsid, as reported for CPV-binding to the canine TfR (53,54) and in the adaptation of CPV to the fox TfR (15). Hence, the acquisition of new host ranges may occur in a step-wide fashion, where one or a small number of mutations will allow initial establishment, as seen here after growth on grey fox cells, and perhaps followed by a small number of additional mutations to optimize the virus:host interaction. Similar dynamics may occur with antibody epitopes and CPV’s evasion of humoral immunity, and single mutations have been shown to affect the binding of monoclonal antibodies (14,22,55).

Taken together, our comparison of the both multi-decade-long global-scale and short-term intra-host evolutionary processes provides a uniquely detailed view of the evolution of an emerging virus in the context of rapid host adaptation, for which high levels of intra- and inter host diversity are not defining features. This sits in contrast to the models that are typically presented for RNA viruses of similar or greater evolutionary capacity due to the low fidelity of their RNA polymerases. We therefore propose that while the waiting time for beneficial mutations in CPV is likely longer than that seen in many RNA viruses, once such mutations appear natural selection may act directly and rapidly fix them in the population, facilitated by relatively rapid virus replication. During the early emergence of CPV in dogs, the opportunities for fitness enhancement were high such that multiple advantageous mutations were successfully fixed in the population. Following, this initial period of host adaptation, the relative proportion of beneficial mutations declined and, combined the pleiotropy inherent in CPV, reduced the fixation rate. Thus, our findings emphasize a need to consider multiple facets of a virus’s biology, pathogenesis, and ecology, and not simply mutation rate, when considering its adaptive capabilities.

## MATERIALS AND METHODS

### Viral genome amplification, library preparation and consensus sequencing

Naturally circulating virus samples had all be obtained directly from infected animals, or as diagnostic samples provided directly from infected animals, and had been stored at Cornell University at −80°C since they were collected. Most were feces, but a few were intestinal tissues. In addition, other tissues were used to investigate any tissue specific mutation in an acutely infected dog (“CPV617”). The origin and collection date of all natural samples are listed in Table 1. DNA was extracted from all samples regardless of material type using the E.Z.N.A. Tissue DNA Kit. For experimentally passaged viruses, prior to DNA extraction, culture supernatant was treated with DNAseI to remove any residual plasmid that may have been carried over from transfection. For all samples sequenced, input CPV genome copy numbers were determined via qPCR as in (56), using primers 5’-AAATGAAACCAGAAACCGTTGAA-3’, 5’-TCCCGCGCTTTGTTTCC-3’ and probe (5’-ACAGTGACGACAGCAC-3’) targeting a conserved region of the NS1 gene. In most cases, 10^6^ – 10^7^ copies of the viral DNA were used to initiate the PCRs and sample inputs below 10^-4^ were not used, ensuring proper amplification with minimal PCR cycling and reduced any sampling effects on the detection of SNVs among samples that were deep sequenced (57). In addition, all PCRs for deep sequenced samples were run in triplicate and pooled before library preparation to minimize individual PCR errors and sampling effects. A region of the genome spanning all major reading frames and portions of the 5’ and 3’ UTRs was amplified (see Fig. S1 for primer sequences and locations) and Q5 High Fidelity DNA Polymerase (New England BioLabs) under the recommended conditions and 25 rounds of amplification. Amplification products were purified with 0.45 x volume AMPure XP beads (Beckman Coulter) and 1ng input DNA used to construct barcoded sequencing libraries with the Nextera XT kit (Illumina). Libraries were multiplexed and sequences were determined using Miseq 2×250 Illumina sequencing. Raw sequencing reads were trimmed using BBDuk (https://jgi.doe.gov/data-and-tools/bbtools/bb-tools-user-guide/) to remove all adaptors and low quality regions from reads. The reads were then merged and mapped to a CPV-2 reference (EU659116) with 2 iterations using Geneious Prime v. 2019.0.4 (58) and the majority consensus sequence for each sample was determined.

### Minor variant calling

For samples that were deep sequenced, additional read processing was performed. Using BBNorm (https://jgi.doe.gov/data-and-tools/bbtools/bb-tools-user-guide/bbnorm-guide/), reads were error-corrected and normalized to target 5000-fold coverage per site. Reads for each sample were re-mapped to their consensus sequence and an additional filtering step (BAMUtil:Filt, (59)) was performed to clip base miscalls at the termini of reads which often result in poor assemblies and SNV call errors. Coverage and site frequencies across the region of interest were determined using the PileupParam & ScanBamParam features in Rsamtools (60), and the most abundant minor variant frequency was called for each position.

To define the accuracy and sensitivity of our ability to detect rare sub-consensus SNVs in viral samples when sequencing the CPV genome, we conducted control sequencing studies of artificially created heterogeneous virus populations. These were made from a CPV-2 plasmid (61) spiked with 25% FPV plasmid, and serially diluted 4 times at a 1:4 ratio (Fig. S2A). An input amount of 0.2ng of mixed template DNA (equivalent to ∼2.5×10^7^ copies of the viral genome) was then prepared by PCR as for all of the natural viral samples. The spiked genomes were easily detected in a dose dependent manner at approximately the proportions that were prepared (Fig. S2B & C).

### Phylogenetic analysis

For the full-genome CPV phylogenies we excluded all recombinant viruses (n = 6 in total) as determined by more than 2 different methods in default RDP4 analysis (39). Evolutionary relationships were among remaining sequences (n = 219) were determined using the maximum likelihood (ML) method available in PhyML (62), employing a general time-reversible (GTR) substitution model, gamma-distributed (Γ) rate variation among sites, and bootstrap resampling (1000 replications). Mutations of interest were catalogued manually and annotated on tree tips using the Microreact web-server (63).

### Experimental passages

To experimentally simulate, *in vitro*, a cross-species transmission event, a plasmid derived A CPV-2b virus (p440, sequence CPV39 in this study) (42) was passaged on domestic cat (*Felis catus*) kidney (NLFK) and gray fox (*Urocyon cinereoargenteus*) lung (FoLu) cells. NLFK cells were maintained in McCoy’s 5A and Liebovitz L15 media with 5% fetal calf serum (FCS) while FoLu cells were maintained in Dulbecco modified Eagle medium (DMEM) 10% FCS and both cell lines were grown at 37°C and 5% CO_2_ atmosphere. A common virus stock was generated by transfection of plasmid DNA into NLFK cell culture using Lipofectamine 2000 (Life Technologies, Carlsbad, CA) according to the manufacturer’s instructions. Stock virus was then used to infect individual flasks of FoLu and NLFK cells run in triplicate. For each virus passage, cells were seeded 8-16 hours prior to infection at (∼1×10^5^ cells/mL) in 12.5 cm^2^ area flasks. Prior to inoculation, growth media was removed and cells were washed twice with sterile PBS. 250µl of virus (from stock or previous passage) was added directly to cells, which were then incubated for 1hr to allow infection, after which time conditioned media was added back to the flask for a final volume of 2.5ml. Infected cells were grown until confluent (4 to 6 days) and supernatant was harvested and viral DNA amplification, deep sequencing and variant calling was performed on stock and passaged viruses as previously described.

## ACKNOWLEDGEMENTS

We thank Wendy Weichert and Renee Anderson for expert technical support.

